# *Bacteroides fragilis* modulates gut microbiome community composition, frontal cortex gene expression, and fecal and frontal cortex metabolites in a mouse model of Alzheimer’s disease pathologies

**DOI:** 10.64898/2026.05.29.728805

**Authors:** Kathryn A. Conn, Olivia Schwartz, Shreya Dikshit, Daisy L. Barroso-Montalvo, David Hattan, Chloe Herman, Colin V. Wood, Emily M. Borsom, J. Gregory Caporaso, Allegra Aron, Emily K. Cope

## Abstract

Alzheimer’s disease (AD) is a neurodegenerative disease and the leading cause of dementia among elderly. Gut microbiome alterations precede pathogenesis and may affect disease outcomes. We evaluated the role of *Bacteroides fragilis* in triple transgenic mice modeling AD pathologies (3xTg-AD) and wild-type controls (WT). Subsets of 3xTg-AD and WT mice were longitudinally treated with *B. fragilis* or sterile vehicle control for five consecutive days at 8 weeks of age and then monthly up to 52-56 weeks. Fecal samples were collected fortnightly from 8 through 52-56 weeks of age. Mice were sacrificed at 8 (baseline), 24 (amyloid-β plaques modeled), and 52-56 (amyloid-β plaques and neurofibrillary tangles modeled) weeks of age. Expression of genes involved in neuroinflammation and neurotransmission were quantified using reverse transcription quantitative polymerase chain reaction (RT-qPCR). Fecal bacterial microbiota were assessed by sequencing the V4 region of the 16S rRNA gene, and microbiome sequence data were analyzed using QIIME 2. Frontal cortex and fecal metabolomes were evaluated using LC-MS/MS. We observed that *B. fragilis* colonized the gut microbiota of 3xTg-AD mice earlier and more consistently than WT mice. 3xTg-AD mice treated with *B. fragilis* demonstrate lower gene expression of *GFAP, SLC1A3,* and *FOXO3* in the frontal cortex. Consistent with this finding, treatment with *B. fragilis* restores levels of amino acid derivatives and neurotransmitters in 3xTg-AD mice to resemble levels in WT mice. These results highlight the role of the gut microbiome in AD-associated neuroinflammation and neurotransmission and the need for future studies to elucidate the mechanisms underlying these changes.

**Importance:** Previous studies in animals modeling Alzheimer’s disease pathologies and humans living with Alzheimer’s disease demonstrate shifts in the gut microbial community composition prior to and concomitant with pathological onset. The bacterial genus, *Bacteroides*, is commonly found differentially abundant in these studies but its effects on disease outcomes are poorly understood. In this study, we explore the effects of chronic exposure to *Bacteroides fragilis* in triple transgenic mice modeling Alzheimer’s disease pathologies and healthy, wild-type controls. We observed changes in microbial community composition in mice modeling Alzheimer’s disease when treated with *B. fragilis,* and associated changes in neuroinflammation, biomarkers of neurotransmission, and the brain metabolome. Taken together, these results suggest that *Bacteroides fragilis* exerts neuromodulatory effects that may be beneficial in Alzheimer’s disease.

## Introduction

Alzheimer’s disease (AD) dementia affects approximately 1:9 Americans over the age of 65^1^. Hallmark pathologies of AD include amyloid-β plaques, neurofibrillary tangles, and neuroinflammation^1,2^. Neuroinflammation is primarily a result of the activation of microglia and astrocytes, immune cells of the brain’s innate immune system^3,4^. Currently, no cure exists for AD, and Phase III clinical trials of amyloid-β-clearing drugs have failed to rescue cognitive deficits^5–7^. Thus, interest has been growing in understanding peripheral pathophysiology which may underlie disease outcomes. Recent studies of AD indicate that shifts in gut microbial communities prior to development of plaque and tangle pathologies may contribute to disease via the “gut-microbiome-brain axis,” although mechanisms explaining these changes have not been established.

Although gut microbial features enriched or depleted in humans living with AD or mouse models vary across studies, some taxa are consistently differential in the AD microbiome. The genera *Bacteroides, Lactobacillus, Bifidobacterium, Turicibacter, Akkermansia, Prevotella, Alistipes,* and *Blautia* are frequently found as differentially abundant in AD^8–11^. We previously demonstrated that *Bacteroides acidifaciens,* part of the *Bacteroides fragilis* group^12^, was enriched in 3xTg-AD mice after the development of amyloid-β plaques^9^. In this study, we demonstrated that the baseline (8 week) gut microbiome is a predictor for mice modeling AD before development of plaque or tangle pathologies^9^. This suggests a role for the microbiome in AD development prior to pathogenesis. A recent study in cognitively normal individuals with biomarkers of preclinical AD demonstrated changes in the gut microbiome associated with biomarkers of tauopathy and amyloidosis^13^. Experimental approaches are needed to understand the role of specific gut microbiota in the gut-microbiome-brain axis.

The effect of *B. fragilis* on neurological outcomes range from positive to negative depending on the model. One study demonstrated that fecal microbiota transplantation (FMT) from human donors with confirmed AD into Thy1-C/EBPβ transgenic mice, which model age-dependent plaques and tangles, results in increased relative abundance of *B. fragilis* compared to mice who received FMT from healthy individuals^14^. This increased abundance was associated with cognitive decline and microglia activation^14^. Other studies have suggested a positive role of *B. fragilis* in the gut-microbiome-brain axis. In a study of mice modeling behaviors associated with autism spectrum disorder (ASD), *B. fragilis* exerted a protective role and corrected defects in ASD-associated behaviors associated with production of the gut metabolite 4-EPS^15^. A recent clinical trial in children with ASD demonstrated improved behavioral outcomes and GI symptoms after consuming *B. fragilis* for 16 weeks^16^. *Bacteroides* produce and consume mammalian neurotransmitters, including the excitatory neurotransmitter glutamate, the inhibitory neurotransmitter ɣ-aminobutyric acid (GABA), and their precursor, glutamine^17,1818–20^. Here, we evaluated the longitudinal effects of orally-administered *B. fragilis* in a model of AD pathologies. We hypothesized that 3xTg-AD mice chronically exposed to *B. fragilis* would demonstrate altered gut microbial communities which shift towards a WT microbiome, reduced neuroinflammatory markers, and restoration of brain and fecal metabolome toward a WT phenotype.

## Materials and Methods

### Genotypes

We used a triple transgenic mouse model of AD (3xTg-AD; B6;129-Tg(APPSwe,tauP301L)1Lfa *Psen1^tm1Mpm^*/Mmjax) which models amyloid-β plaques (APP[Swe], PSEN1[M146V]) and tau neurofibrillary tangles (MAPT[P301L], Jackson Laboratory). Wild type mice of the same genetic background were used as controls (B6129SF2/J, Jackson Laboratory). Genotyping was performed with PCR to confirm the presence of the transgenes (Supplemental methods). Experimental mice were bred in-house at the Biological Sciences Vivarium at Northern Arizona University. Female mice were selected for this study. This study was approved by Northern Arizona University’s Institutional Animal Care and Use Committee (IACUC protocol number 21-019). All experiments were performed following the ARRIVE guidelines and regulations^21^.

### Mouse colonies

Mice were assigned to age groups (8 weeks [baseline], 24 weeks [amyloidosis], and 52-56 weeks [amyloidosis and tauopathy]) and *B. fragilis* or vehicle control treatment groups. A total of 111 mice were used in this study. Detailed methods on mouse colonies can be found in Supplemental methods.

### Treatment preparation of *Bacteroides fragilis*

*B. fragilis* ATCC 25285 was purchased from the American Type Culture Collection (ATCC, Manassas, Virginia). We followed the protocol established by Hsiao et. al. for treatment preparation with the only modification being 1×10^9^ CFU/mL of *B. fragilis* in our dosages^15^. Preparation of *B. fragilis* doses are described in the Supplemental Methods.

### Bacteroides fragilis treatment

Dosages of *B. fragilis* or sterile vehicle control were mixed into 4mL of sugar-free applesauce. We administered 1.25mL of treatment/applesauce mixture per mouse in each cage spread over standard food pellets. At eight weeks of age, mice received 5 consecutive days of treatment with *B. fragilis* or vehicle control and then once monthly thereafter (Fig. 1).

**Figure 1.**
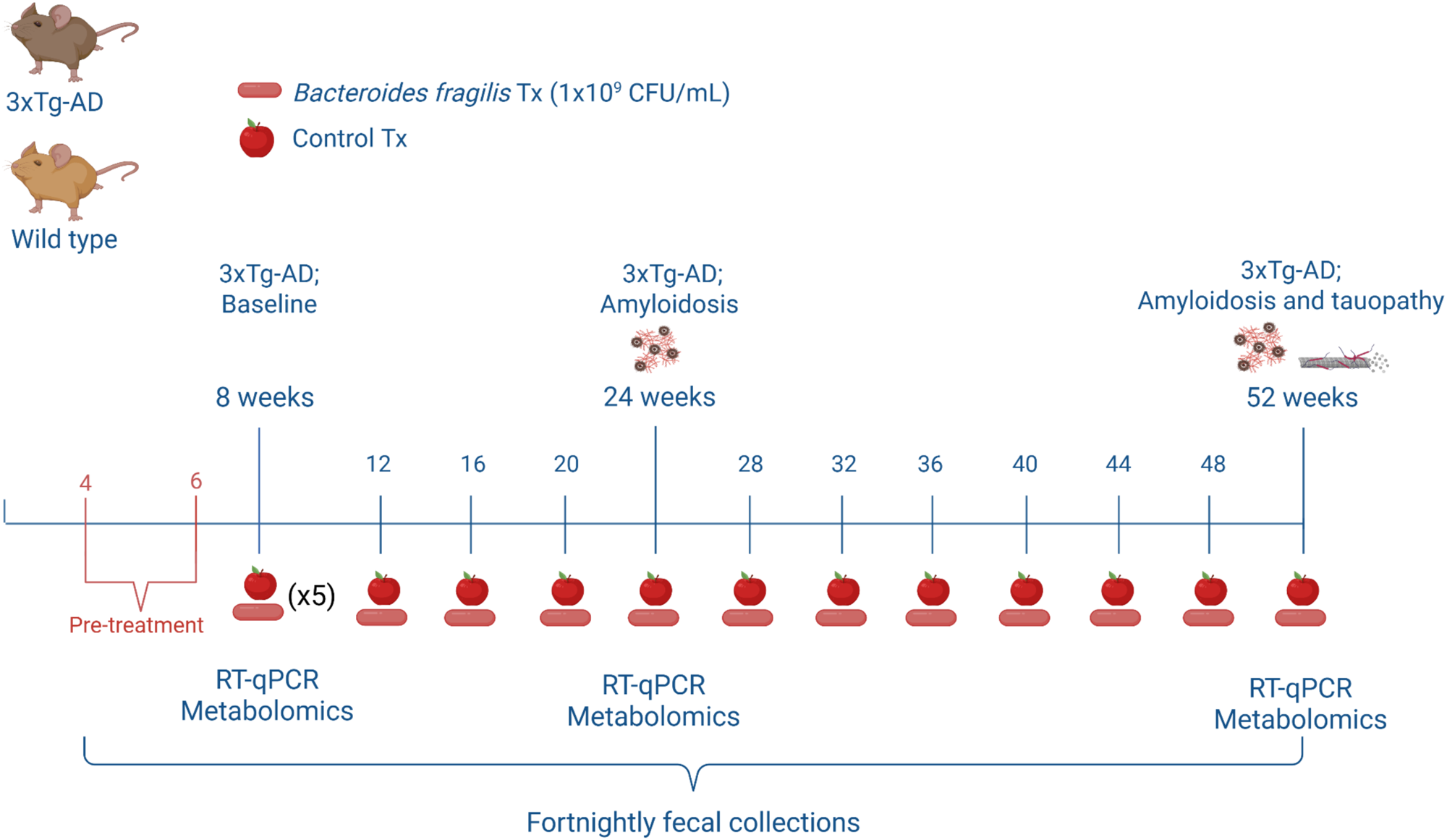
Experimental design and timeline for treatment with *B. fragilis* and disease pathological onset. Fecal samples were collected fortnightly beginning at four weeks of age. Treatment with *8. fragilis* or sterile vehicle control began at eight weeks of age for five consecutive days and then once every four weeks following. 1ml treatments of *8. fragi/is* or vehicle control were mixed with 4ml of unsweetened applesauce and spread over food pellets in the cage. Dissections and tissue harvesting was performed at eight weeks of age (baseline), 24 weeks of age (amyloidosis), and 52 weeks of age (amyloidosis and tauopathy). Image created using <u>BioRender.com</u>

### Dense longitudinal fecal microbiome sampling

Fresh fecal samples were collected from each mouse fortnightly beginning at four weeks of age through their terminal timepoint. Fecal samples were collected by temporarily restraining mice by holding the base of their tails and collecting one to two fecal pellets in 1mL of 1X sterile phosphate buffered saline (PBS). Fecal samples were immediately transferred to a -70℃ freezer until further processing.

### Sample collection for metabolomics

Frontal cortex and fecal samples were collected for metabolomic analysis during dissection. The brain was washed with cold PBS and then the frontal cortex was gently separated from the brain using a sterile scalpel. Brain and fecal samples were immediately placed on ice until they were transferred to permanent storage in a -70℃ freezer.

### Nucleic Acid Extraction

DNA and RNA were extracted from frontal cortex samples using a MagMAX™ Pathogen RNA/DNA Kit (Thermo Fisher Scientific). RNA was purified using a Thermo Scientific™ RapidOut DNA Removal Kit (Thermo Fisher Scientific). 3μL (69-216ng) of RNA per sample were reverse transcribed into complementary DNA (cDNA) using an RT^2^ First Strand Kit (Qiagen, Redwood City, California). Nucleic acid extraction details can be found in Supplemental Methods.

### Microbiome sequencing

We sequenced the V4 region of the 16S rRNA gene on the Illumina MiSeq as previously described^9^. PCR conditions and sequencing library preparation are fully described in the Supplemental Methods.

### Reverse Transcriptase Quantitative PCR

Frontal cortex cDNA was quantified on a NanoDrop 2000 (Thermo Fisher Scientific) and standardized to 10ng/μL for RT-qPCR. Samples were run in duplicate and an average of the cycle threshold was calculated for each sample. A custom RT-qPCR panel included nine genes of interest and four housekeeping genes (Table 1; Qiagen GeneGlobe). We verified that all samples were free of genomic DNA through an independent PCR assay using primers for the mouse ribosomal protein L13A (RPL13A). 72 frontal cortex samples were used (Table 2).

**Table 1.** RT-qPCR assay genes and their function in GABA and glutamate transport and neuroinflammation.

| **Gene** | **Description** |
| --- | --- |
| *GABRA1* | GABA transport |
| *GABRA2* | GABA transport |
| *GABRA3* | GABA transport |
| *SLC6A12* | Glutamate transport |
| *SLC1A3* | Glutamate transport |
| *FOXO3* | Neuroinflammation |
| *MRC1* | Endocytosis of glycoproteins via macrophages |
| *GFAP* | Glial cell, neuron, BBB support |
| *IL6* | Inflammation |
| *GAPDH* | Housekeeping |
| GDC | Genomic DNA Control |
| PPC | Positive Control |
| RTC | Reverse Transcriptase Control |

**Table 2.** Treatment groups sizes.

|  | **WT, Bf** | **WT, Ctrl** | **3xTg-AD, Bf** | **3xTg-AD, Ctrl** |
| --- | --- | --- | --- | --- |
| **8wks** | n = 10 | n = 10 | n = 10 | n = 8 |
| **24wks** | n = 10 | n = 10 | n = 9 | n = 8 |
| **52wks** | n = 9 | n = 8 | n = 9 | n = 10 |

### Microbiome bioinformatics analysis

All bioinformatic analyses were performed with QIIME 2 2024.10.1^22^. All analyses are detailed in the Supplemental Methods and all commands used are included in the QIIME 2 Provenance Replay.

### Fecal sample extraction for metabolomics

Tissue samples were frozen at -80°C upon receipt at the University of Denver. Further details are described in Supplemental Methods.

### Frontal cortex sample extraction for metabolomics

Extraction was performed as described in literature^23^. Further details are described in Supplemental Methods.

### RP-LC-MS/MS analysis

2µL of samples were injected into a Vanquish UHPLC system coupled to a Q-Exactive orbitrap mass spectrometer (Thermo Fisher Scientific). A nonpolar C18 porous core column (Kinetex Core-shell polar C18, 150 x 2mm, 2.6µm particle size, 100 Å pore size, Phenomenex, Torrance, USA) was used for chromatographic separation. Details on gradient and instrument parameters are included in Supplemental Methods.

### Hydrophilic lipophilic interaction chromatography (HILIC) tandem MS/MS analysis

10μL of frontal cortex and 5μL of fecal samples were injected into a Vanquish UHPLC system coupled to a Q-Exactive orbitrap mass spectrometer (Thermo Fisher Scientific). An amide column (XBridge BEH Amide Column, 130 Å pore size, 1.7µm particle size, 2.1mm X 100m, Waters, Milfrod, MA). Chromatographic separation was conducted with the reagents and gradient described by Lai, et al^24^. Other parameters used are detailed in Supplemental Methods.

### Feature detection of RP-LC-MS/MS

mzmine 4.3.0^25^ was utilized to perform all feature finding steps. MS/MS spectra were converted to .mzML files using msConvert (ProteoWizard)^26^. MS1 feature extraction and MS/MS pairing were performed with mzmine 4.3.0. Parameters used are detailed in Supplemental Methods.

### Feature detection of HILIC-MS/MS

mzmine 4.3.0^27^ was utilized to perform all feature finding steps. MS/MS spectra were converted to .mzML files using msConvert (ProteoWizard)^26^. MS1 feature extraction and MS/MS pairing were performed with mzmine 4.3.0. Parameters used are detailed in Supplemental Methods.

### Statistical analysis

Analysis of metabolomics data was performed in R Studio version 4.4.1. Quality of LC-MS/MS runs was assessed using coefficient of variance (CV) values for internal standards (Supplemental Table S1-S3) and for quality control mixes analyzed throughout each run (Supplemental Table S4-S6). Batch effect (Supplemental Fig. 1-3) was corrected as described in Supplemental Methods. Annotation of metabolites are described in Supplemental Methods (Supplemental Fig. 4). All code can be found at https://zenodo.org/records/17822090?token=eyJhbGciOiJIUzUxMiJ9.eyJpZCI6Ijk1NWIwMTA4LThiMDUtNDQxZS04ZTE1LTNiMWJiMGI3NjdlMiIsImRhdGEiOnt9LCJyYW5kb20iOiIxYzQyY2RmNGM3ZTU1NmE4MDcwNzg2YmJhNzQyYTliZSJ9.8FQbzo_Zg4OLth_ONQqtt8evmbP6zpFsw9g9-1Q-nM7hTkloNEvzh-owRnhx4u4gq7ooJhACJWn-j5iGa02tsQ

Analysis of RT-qPCR data and creation of violin plots were performed using PRISM (GraphPad version 10.1.1, GraphPad Software, Boston, Massachusetts). Fold changes for frontal cortex RT-qPCR analysis were performed using the cycle threshold (2^-ΔΔCT^) method. Fold change for each biomarker was calculated relative to *GAPDH*. Group analyses were performed using a Mann-Whitney test and then all p-values were corrected for multiple comparisons using Holm-Bonferroni^28^.

### Annotation of metabolites

Samples were analyzed alongside a set of 600 standards (IROA Mass Spectrometry Metabolite Library Standards) using HILIC and RP-LC to achieve level 1 annotations of metabolites of interest^29^, where features were annotated by matching retention time, m/z, and MS2 fragmentation to standards. Further details, and other annotation methods are described in Supplemental Methods.

### Data Availability

Metabolomics raw and processed data can be found with the following MASSIVE IDs: Fecal RP-LC-MS/MS (<u>MSV000095774</u>), fecal HILIC-MS/MS (<u>MSV000097845</u>), and frontal cortex HILIC-MS/MS (<u>MSV000098035</u>). Raw microbiome sequence data were deposited in NIH Sequence Read Archive (SRA) accession number: PRJNA1417316.

## Results

### Treatment with *Bacteroides fragilis* downregulates glial gene expression in the frontal cortex of mice genetically predisposed to AD pathologies

RT-qPCR was performed on frontal cortex RNA to determine the effects of mouse genotype, age, and chronic exposure to *B. fragilis* on biomarkers of neurotransmitter receptors and inflammation (Table 1). We demonstrate a significant increase of glial fibrillary acidic protein (*GFAP*) gene expression in 3xTg-AD mice aged to 52-56 weeks compared to 8 weeks within the vehicle control treatment group (Mann-Whitney, Holm-Bonferroni p=0.018). In 3xTg-AD mice treated with *B. fragilis*, *GFAP* gene expression at 52-56 weeks compared to 8 weeks of age, more resembled what we observed in the WT mice (Mann-Whitney, Holm-Bonferroni p=0.8970, Fig. 2A). We observed a decrease in expression of the transcription factor forkhead box O3 (*FOXO3*) gene in 52-56 week old 3xTg-AD mice given *B. fragilis* compared to 52-56 week 3xTg-AD mice given vehicle control (Mann-Whitney, Holm-Bonferroni p=0.009; Fig. 2B). Consistent with this observation, *SLC1A3*, a gene which encodes the protein excitatory amino acid transporter 1 ([*EAAT1*] Mann-Whitney, Holm-Bonferroni p=0.009, Fig. 2C) was also decreased at the 52-56 week timepoint in 3xTg-AD mice given *B. fragilis* compared to 52-56 week mice given vehicle control. Because *B. fragilis* is a potent producer of GABA in the gut, we next sought to analyze levels of the GABA_A_ receptor subunits *GABRA1*, *GABRA2*, and *GABRA3.* After correction for multiple comparisons, we observed no change across genotype, age, or treatment types. (Fig. 2D-F). We also analyzed expressions of *SLC6A12*, *MRC1*, and *IL6*. None of these genes were significantly different across groups (Supplemental Fig. 5).

**Figure 2.**
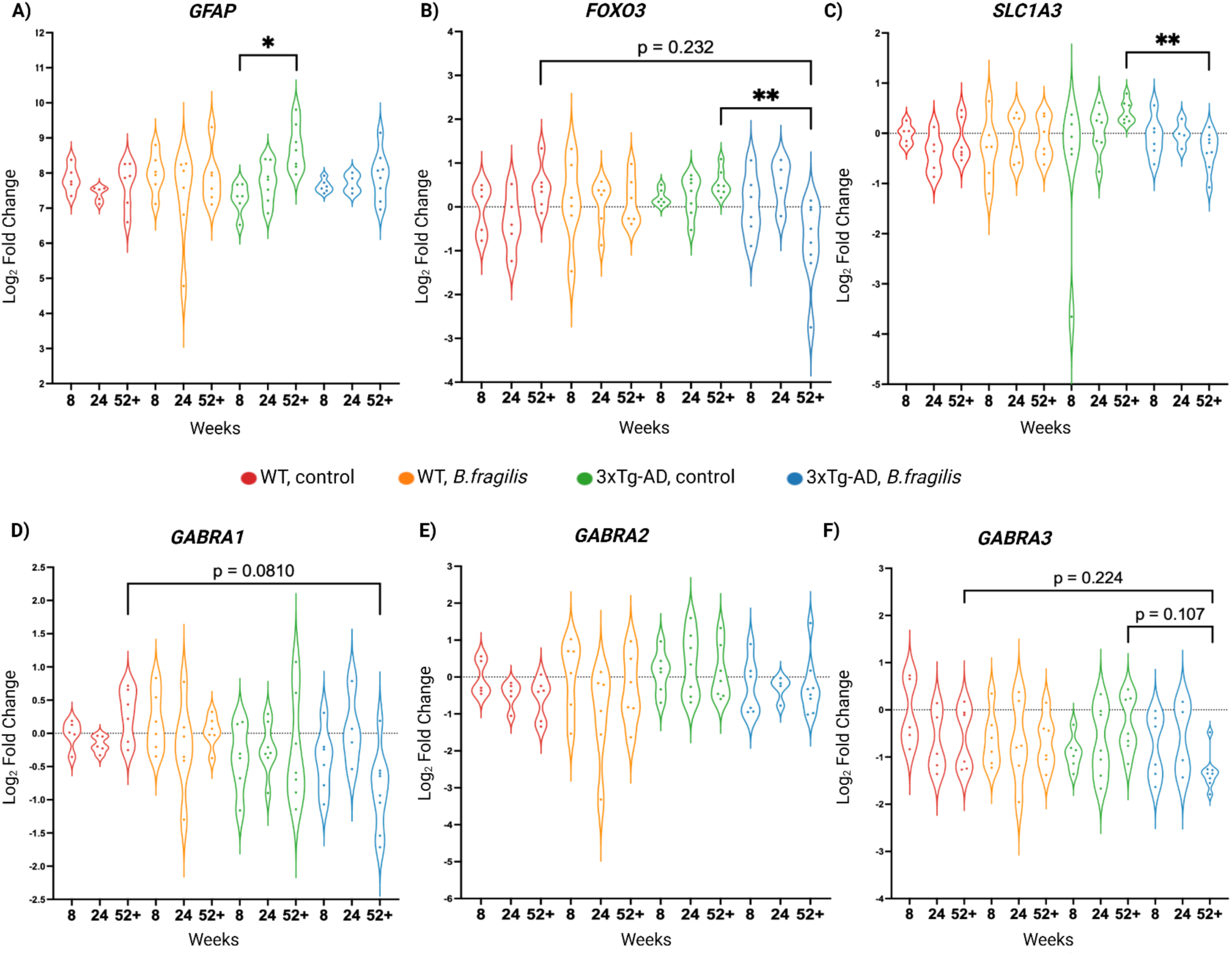
Log2 fold changes of frontal cortex mRNA expression (normalized to *GAPDH).* A) Frontal cortex mRNA expression of *GFAP* relative to GAPDH (n = 4-8, Welch’s ANOVA p=0.034). mRNA expression of *GFAP* is upregulated between 8 and 52-56 weeks of age in 3xTg-AD treated with sterile vehicle control (Holm-Bonferroni p=0.0360). B) Frontal cortex mRNA expression of *FOX03* relative to *GAPDH(n* = 4-8, Welch’s **AN**OVA p=0.131). mRNA expression of *FOX03* is significantly downregulated in 3xTg-AD mice treated with *B. fragilis* at 52-56 weeks of age relative to 3xTg-AD mice treated with sterile vehicle control at 52 weeks of age (Holm-Bonferroni p=0.009). C) Frontal cortex mRNA expression of *SLCJA3* relative to *GAPDH(n* = 4-8, Welch’s ANOVA p=0.014). mRNA expression of *SLCJA3* is significantly downregulated in 3xTg-AD mice treated with *B.fragilis* at 52-56 weeks of age relative to 3xTg-AD mice treated with sterile vehicle control at 52 weeks of age (Holm-Bonferroni p=0.009). D) Frontal cortex mRNA expression of GABRA1 relative to *GAPDH(n=4-8,* Welch’s ANOVA p=0.070). mRNA expression of *GABRA1* is not significantly decreased across age, genotype, or treatment type. E) Frontal cortex mRNA expression of *GABRA2* relative to *GAPDH* (n=4-8, Welch’s ANOVA p=0.432). mRNA expression of *GABRA2* is not significantly decreased across age, genotype, or treatment type. F) Frontal cortex mRNA expression of *GABRA3* relative to *GAPDH* (n=4-8, Welch’s ANOVA p=0.101). mRNA expression of *GABRA3* is not significantly decreased across age, genotype, or treatment type. *ps0.05, ** ps0.01

We next wanted to understand if expression of *SLC1A3,* a glutamate transporter, and FOXO3 were correlated, as FOXO3 is correlated with glutamine synthetase. Linear models were fit to the following timepoints; baseline, amyloidosis, amyloidosis and tauopathy, all samples, and “adult” (includes both ‘amyloidosis’ and ‘amyloidosis and tauopathy’ timepoints). We demonstrate for the first time that mRNA expression of *FOXO3* and *SLC1A3* are correlated, although this is not affected by mouse genotype nor treatment with *B. fragilis* (Fig. 3A-E).

**Figure 3.**
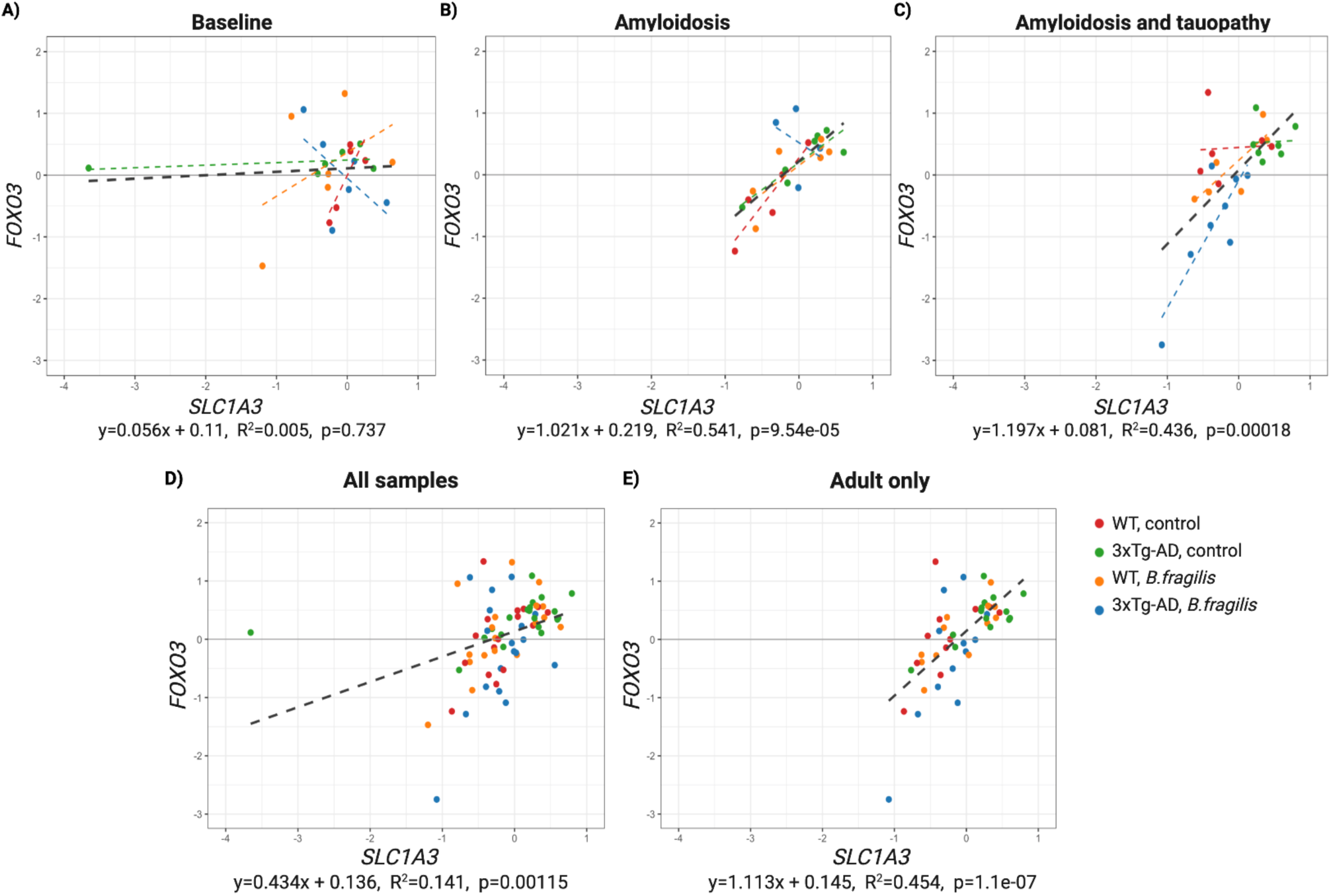
Linear regression of frontal cortex log2mRNA expression of *FOX03* and *SLC1A3.* Samples are colored by group (genotype and treatment type) and linear regressions are fitted to each group. Black lines give the linear regression for all samples in the plot. Colored lines give the linear regression for the corresponding group. A) mRNA expression of *FOX03* and *SLC7A3* are not correlated at the baseline time point (R^2^=0.005, p=0.737). mRNA expression grouped by genotype and treatment type are not significantly correlated at the baseline time point (WT, control: Holm-Bonferroni; p=0.353; WT, *B.fragilis:* Holm-Bonferroni; p=0.723; 3xTg-AD, control: Holm-Bonferroni; p=0.723; 3xTg-AD, *8.fragilis:* Holm-Bonferroni; p=0.637). B) mRNA expression of *FOX03* and *SLC1A3* are correlated at the amyloidosis time point (R^2^=0.541, p=9.54e-05). mRNA expression grouped by genotype and treatment type at the amyloidosis time point is significantly correlated in 3xTg-AD mice treated with sterile vehicle control (3xTg-AD, control: Holm-Bonferroni; p=0.036) but not in 3xTg-AD mice treated with *B. fragilis* or WT mice in either treatment group (WT, control: Holm-Bonferroni; p=0.093; WT, *8. fragilis:* Holm-Bonferroni; p=0.1221; 3xTg-AD, *8. fragilis:* Holm-Bonferroni; p=0.655). C) mRNA expression of *FOX03* and *SLC7A3* are significantly correlated at the amyloidosis and tauopathy time point (R^2^=0.436, p=0.00018). *FOX03* and *SLC7A3* are significantly correlated in 3xTg-AD mice treated with *8.fragilis* at the amyloidosis and tauopathy time point (Holm-Bonferroni; p=0.038) but not in 3xTg-AD mice treated with sterile vehicle control or WT mice in either treatment group (WT, ctrl: Holm-Bonferroni; p=1.000, WT, *B.fragilis.* Holm-Bonferroni; p=0.165, 3xTg-AD, ctrl: Holm-Bonferroni; p=1.000). D) mRNA expression of *FOX03* and *SLC7 A3* genes are significantly correlated when all samples are included (R^2^=0.141, p=0.001). E) mRNA expression of *FOX03* and *SLC7A3* genes are significantly correlated in adult mice (excluding mice at the baseline time point; R^2^=0.454, p=1.1e-07).

### Reversal of elevated amino acid and neurotransmitter levels in frontal cortex of 3xTg-AD mice treated with *Bacteroides fragilis*

HILIC-MS/MS was performed on a subset of frontal cortex samples to assess changes in the polar metabolome. Differences in the polar frontal cortex metabolome composition at 8-, 24-, and 52-56 weeks of age were assessed using PCA (Principal Component Analysis) of the robust center log ratio (RCLR) transformed extracted peak areas for the polar metabolome. While separation by both genotype and treatment can be observed visually for all weeks, sample numbers were too low to measure any significance (Supplemental Fig. 6). To assess whether mouse genotype and chronic exposure to *B. fragilis* altered individual frontal cortex metabolites at 52-56 weeks of age, a Kruskal-Wallis test was performed on each of the features detected and 38 features were found to be significant (p<0.05, Supplemental Table S7-S8). Of the significant features, many *in silico-*predicted amino acid derivatives and aminosaccharide derivatives exhibited a decrease in abundance from WT to 3xTg-AD mice treated with vehicle control^30^ (Fig. 4, Supplemental Table S8, Supplemental Fig. 7); treatment of 3xTg-AD genotype with *B. fragilis* increased levels of these metabolites toward that observed in the WT (Fig. 4B, Supplemental Fig. 7)^31,32^. We found that level 1 annotated amino acids (including serine, tyrosine, methionine, and glutamine) along with neurotransmitters and neurotransmitter precursors GABA and choline exhibited a similar trend,^33^ though these changes are non-significant (Fig. 4B, Supplemental Fig. 8)^31,32^.

**Figure 4.**
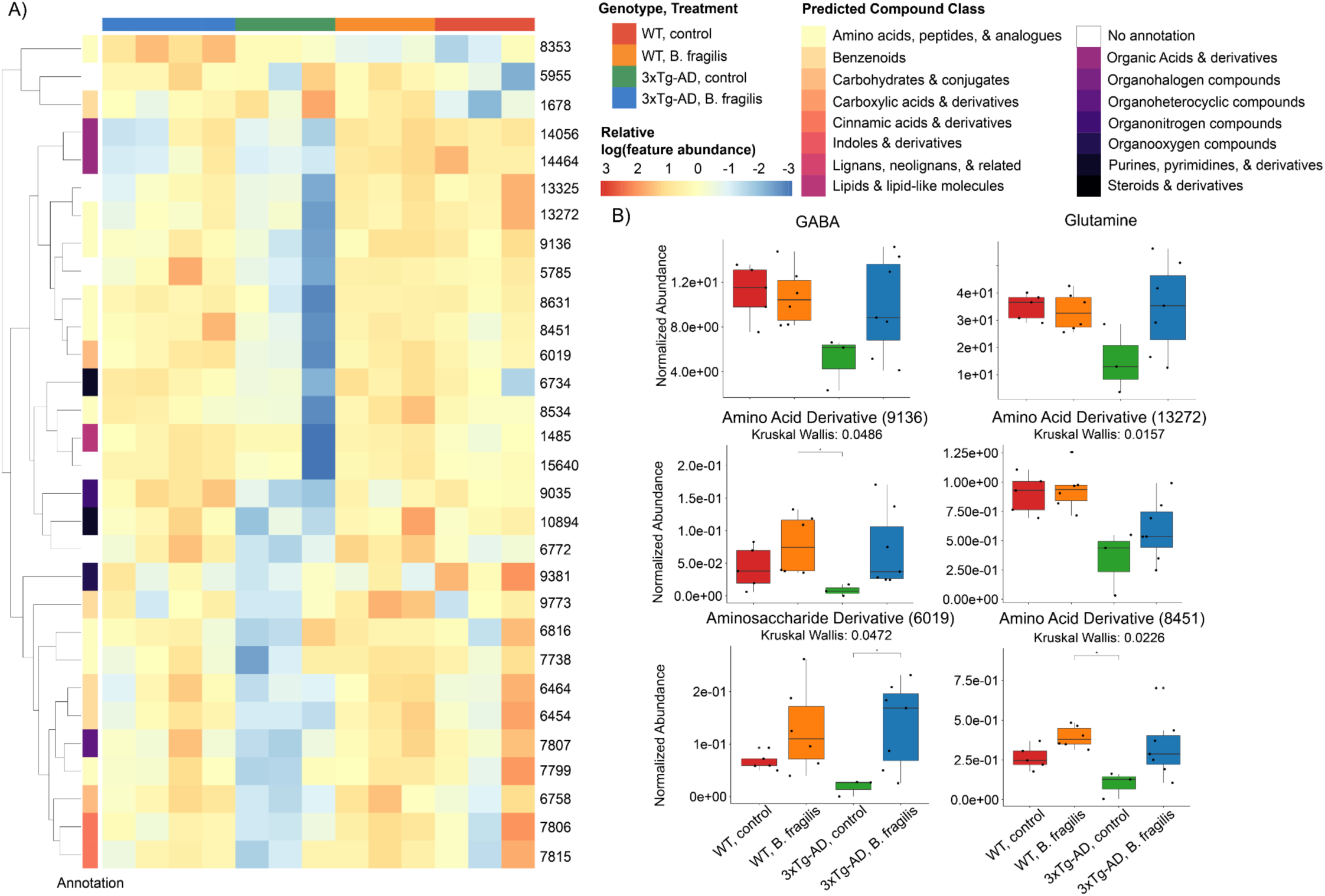
Changes in the frontal cortex (FC) polar metabolome at week 52-56 analyzed using HILIC-MS/MS. **A)** Heatmap of significant FC features at week 52-56. Each sample is colored based on genotype and treatment type (top bar) and each in silico predicted compound class is differentiated by color on a rainbow scale (side bar). The intensity of color shown in the heatmap is based upon the relative log(feature abundance) for each feature. Each feature ID is indicated on the right side of the heatmap plot. B) Boxplots of a subset of features at 52-56 weeks of age. GABA and glutamine are level 1 matches to authentic standards, while feature IDs 9136, 13272, 6019, and 8451 were annotated via *in silica* annotation tools by putative molecular classes. Significance with Kruskal-Wallis p-values for each metabolite are listed, and* indicates p<0.05 in post-hoc Dunn’s analysis with Benjamini-Hochberg (BH) correction.

### *Bacteroides fragilis* is detected in the GI tract of 3xTg-AD mice earlier and more consistently than WT mice

To assess longitudinal changes in the gut microbiome composition and the effects of chronic exposure to *B. fragilis* on bacterial communities, we performed 16S rRNA gene sequencing of the V4 region of bacterial DNA. We collected samples from each mouse fortnightly from 4 weeks of age through 52-56 weeks of age (n=111 mice and n=999 samples, Table 3). A total of 14,836,617 sequence reads were generated with a range of 1,435 to 146,123 reads per sample and a mean of 14,851.5 reads resulting in 2,776 unique amplicon sequence variants (ASVs). For downstream analysis of alpha and beta diversity, samples underwent 100 iterations of rarefaction to a sampling depth of 5,000 sequences/sample, followed by the computation of averaged diversity metrics across the 100 iterations. As a result, 302 samples were excluded from analysis.

**Table 3.** Treatment groups sizes for age, genotype, and treatment with B. fragilis or vehicle control.

|  | **WT, Ctrl** | **WT, B. fragilis** | **3xTg-AD, Ctrl** | **3xTg-AD, B. fragilis** |
| --- | --- | --- | --- | --- |
| **8wks** | n = 10 | n = 10 | n = 8 | n = 10 |
| **24wks** | n = 10 | n = 10 | n = 8 | n = 9 |
| **52wks** | n = 8 | n = 9 | n = 10 | n = 9 |

Longitudinal volatility analysis demonstrated that *B. fragilis* was detectable in 3xTg-AD mice immediately following the initial 5-day exposure and persisted throughout the study period (Fig. 5A). However, *B. fragilis was not consistently detected in WT* mice until approximately 24-26 weeks of age. This pattern corresponds with observed native *Bacteroides* species in the WT fecal microbiome from the earliest timepoints but these taxa were not detected in 3xTg-AD mice (Fig. 5B).

**Figure 5.**
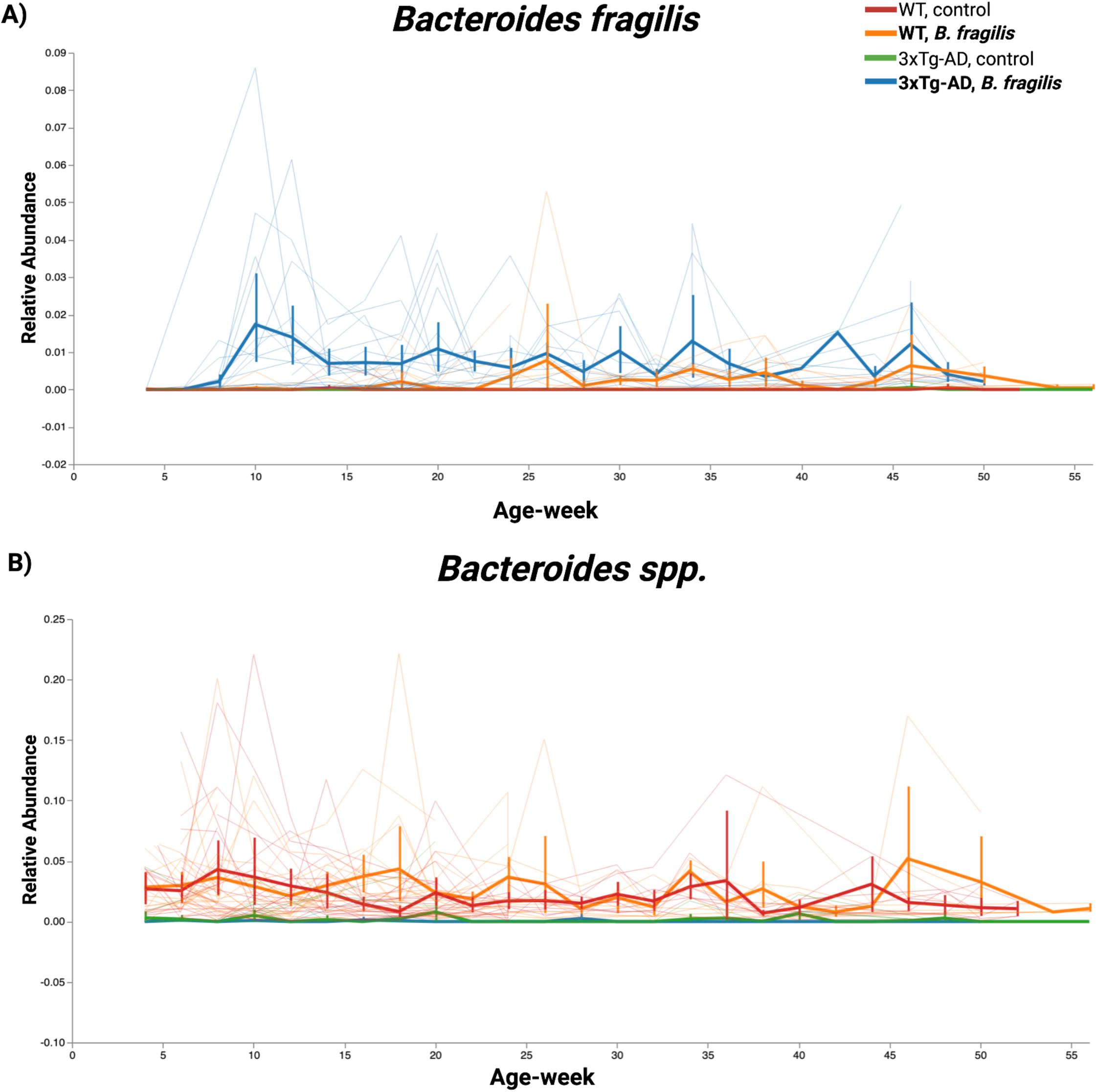
Volatility plots of the relative abundance of *Bacteroides.* Thin spaghetti lines represent longitudinal feature volatility from a single mouse and thick lines are the average of the feature observed in each group. A) Feature volatility of *Bacteroides fragilis* sampled fortnightly from 4-52 weeks of age. B) Feature volatility of *Bacteroides spp.* sampled fortnightly from 4-52 weeks of age.

### Genotype then treatment drives fecal bacterial microbiome and metabolome

We used the Shannon Diversity Index to measure quantitative differences and Faith’s Phylogenetic Diversity (PD) to measure qualitative differences in alpha diversity. Alpha diversity was assessed at baseline, amyloidosis, and amyloidosis and tauopathy timepoints. No significant differences were observed in Shannon Diversity nor Faith’s PD across genotype nor treatment at the baseline (Kruskal-Wallis, Shannon p=0.081, Faith’s PD p=0.758) or amyloidosis (Kruskal-Wallis, Shannon p=0.294, Faith’s PD p=0.201). Shannon diversity was not significantly different at amyloidosis and tauopathy timepoint (Kruskal-Wallis, Shannon p=0.194, Supplemental Fig. 9A-C). However, a significant difference was observed using Faith’s PD across all groups at the amyloidosis and tauopathy timepoint (Kruskal-Wallis; 0.039, Supplemental Fig. 9D-F), but pairwise comparisons showed no significant difference across groups after correction for multiple comparisons (Supplemental Table 9).

We show that genotype is the primary factor driving dissimilarity using Unweighted UniFrac distances (Table 4). Treatment with *B. fragilis* alters the microbiome composition within genotype at all timepoints (Table 4, Supplemental Fig. 10). Volatility analysis of Unweighted UniFrac (which considers phylogenetic relatedness but not ASV abundance) of Axis 1, the most explanatory variable, shows that the gut microbiomes of 3xTg-AD and WT mice remain distinct across the study period (Fig. 6A). In contrast, Axis 2 shows that the gut microbiomes have modest compositional differences between genotypes which converge with age, and Axis 3 shows no difference across genotype nor treatment type (Fig. 6B-C). These changes were supported by volatility plots of individual taxa which showed distinct separation across genotypes and, in some cases, treatment type (Fig 6D-G). Volatility analysis of Weighted UniFrac (which considers phylogenetic relatedness and abundance of an ASV) generally show that the gut microbiomes of 3xTg-AD and WT mice are compositionally similar when weighting high abundance features throughout their lives regardless of treatment type (Supplemental Fig. 11A-C). Pairwise comparisons at the amyloidosis timepoint showed significant differences between 3xTg-AD given *B. fragilis* and WT mice given vehicle control (PERMANOVA; p=0.024, q=0.048, pseudo-F=3.698), between 3xTg-AD given vehicle control and WT mice given vehicle control (PERMANOVA; p=0.004, q=0.015, pseudo-F=4.845), and between 3xTg-AD given vehicle control and WT given *B. fragilis* (PERMANOVA; p=0.005, q=0.015, pseudo-F=4.854, Table 5). Pairwise comparisons at the amyloidosis and tauopathy timepoint showed significant differences between 3xTg-AD given control and WT given *B. fragilis* (PERMANOVA; p=0.008, q=0.048, pseudo-F=3.948, Table 5).

**Figure 6.**
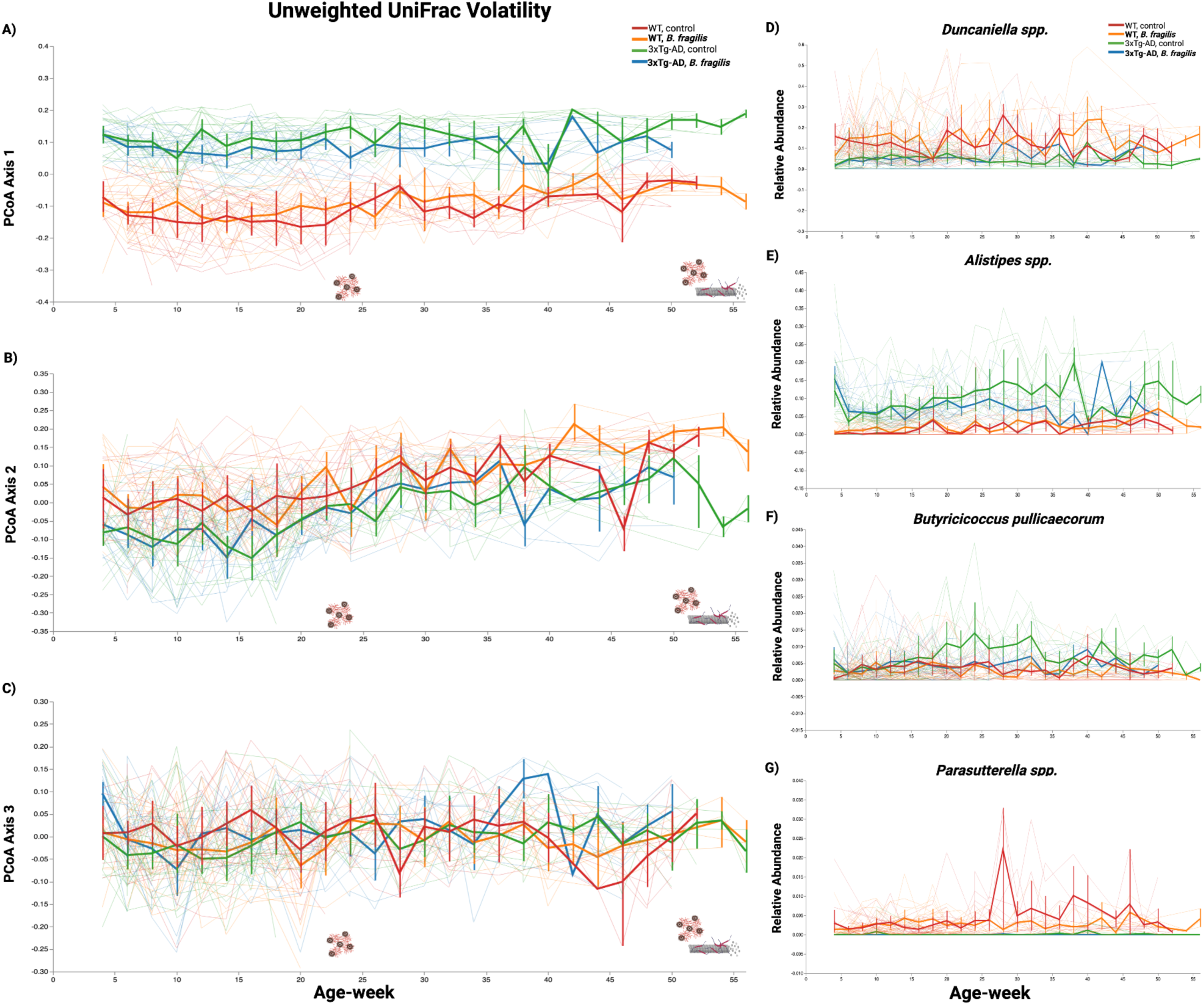
Unweighted UniFrac and individual feature volatility. A) PCoA Axis 1 Unweighted UniFrac. Gut microbiomes remain compositionally distinct between 3xTg-AD and WT mice. B) PCoA Axis 2 Unweighted UniFrac. Gut microbiomes show modest compositional differences between 3xTg-AD and WT mice. C) PCoA Axis 3 Unweighted UniFrac. Gut microbiomes are compositionally indistinguishable between 3xTg-AD and WT mice. D) Feature volatility plot of *Duncaniella spp.* 3xTg-AD mice are depleted of *Duncaniella* in their early lives relative to WT mice. E) Feature volatility plot of *Alistipes spp.* 3xTg-AD mice demonstrate a greater relative abundance of *Alistipes* relative to WT mice in their early lives and may converge with age. F) Feature volatility plot of *Butyricicoccus pul/icaecorum.* 3xTg-AD mice treated with vehicle control demonstrate a greater relative abundance of *Butyricoccus pullicaecorum* beginning when the mice have been characterized to begin modeling amyloid-13 plaques and converging with 3xTg-AD mice treated with *B. fragilis* and both WT groups around the onset of the neurofibrillary tangle (tauopathy) pathology. G) Feature volatility plot of *Parasutterella sp.* 3xTg-AD mice in both treatment groups are depleted of *Parasutterella* relative to WT mice.

**Table 4.** Pairwise PERMANOVA comparisons of Unweighted UniFrac across strain and treatment type at ‘baseline,’ ‘amyloidosis,’ and ‘amyloidosis and tauopathy’ time points.

| Unweighted UniFrac pairwise comparisons - baseline |  |  |  |  |  |  |
| --- | --- | --- | --- | --- | --- | --- |
| Group 1 | Group 2 | Sample size | Permutations | pseudo-F | p-value | q-value |
| 3xTg-AD-Bf | 3xTg-AD-ctrl | 44 | 999 | 1.22852 | 0.202 | 0.202 |
|  | WT-Bf | 43 | 999 | 4.180435 | 0.001 | 0.0015 |
|  | WT-ctrl | 38 | 999 | 5.36558 | 0.001 | 0.0015 |
| 3xTg-AD-ctrl | WT-Bf | 45 | 999 | 5.539058 | 0.001 | 0.0015 |
|  | WT-ctrl | 40 | 999 | 6.687382 | 0.001 | 0.0015 |
| WT-Bf | WT-ctrl | 39 | 999 | 1.722233 | 0.035 | 0.042 |
| Unweighted UniFrac pairwise comparisons - amyloidosis |  |  |  |  |  |  |
| Group 1 | Group 2 | Sample size | Permutations | pseudo-F | p-value | q-value |
| 3xTg-AD-Bf | 3xTg-AD-ctrl | 33 | 999 | 2.137444 | 0.005 | 0.006 |
|  | WT-Bf | 35 | 999 | 4.878644 | 0.001 | 0.0015 |
|  | WT-ctrl | 37 | 999 | 5.675205 | 0.001 | 0.0015 |
| 3xTg-AD-ctrl | WT-Bf | 30 | 999 | 5.725373 | 0.001 | 0.0015 |
|  | WT-ctrl | 32 | 999 | 6.106957 | 0.001 | 0.0015 |
| WT-Bf | WT-ctrl | 34 | 999 | 1.692877 | 0.029 | 0.029 |
| Unweighted UniFrac pairwise comparisons - amyloidosis and tauopathy |  |  |  |  |  |  |
| Group 1 | Group 2 | Sample size | Permutations | pseudo-F | p-value | q-value |
| 3xTg-AD-Bf | 3xTg-AD-ctrl | 19 | 999 | 1.614409 | 0.037 | 0.037 |
|  | WT-Bf | 17 | 999 | 3.356835 | 0.001 | 0.002 |
|  | WT-ctrl | 16 | 999 | 2.224605 | 0.001 | 0.002 |
| 3xTg-AD-ctrl | WT-Bf | 18 | 999 | 3.613553 | 0.001 | 0.002 |
|  | WT-ctrl | 17 | 999 | 2.127482 | 0.007 | 0.0105 |
| WT-Bf | WT-ctrl | 15 | 999 | 1.949861 | 0.018 | 0.0216 |

**Table 5.** Pairwise permanova comparisons of Weighted UniFrac across genotype and treatment type at ‘baseline,’ ‘amyloidosis,’ and ‘amyloidosis and tauopathy’ timepoints.

| Weighted UniFrac pairwise comparisons - baseline |  |  |  |  |  |  |
| --- | --- | --- | --- | --- | --- | --- |
| Group 1 | Group 2 | Sample size | Permutations | pseudo-F | p-value | q-value |
| 3xTg-AD-Bf | 3xTg-AD-ctrl | 44 | 999 | 0.241348 | 0.901 | 0.901 |
|  | WT-Bf | 43 | 999 | 1.545059 | 0.203 | 0.2892 |
|  | WT-ctrl | 38 | 999 | 3.30008 | 0.03 | 0.09 |
| 3xTg-AD-ctrl | WT-Bf | 45 | 999 | 1.286979 | 0.241 | 0.2892 |
|  | WT-ctrl | 40 | 999 | 3.062067 | 0.03 | 0.09 |
| WT-Bf | WT-ctrl | 39 | 999 | 1.723154 | 0.185 | 0.2892 |
| Weighted UniFrac pairwise comparisons - amyloidosis |  |  |  |  |  |  |
| Group 1 | Group 2 | Sample size | Permutations | pseudo-F | p-value | q-value |
| 3xTg-AD-Bf | 3xTg-AD-ctrl | 33 | 999 | 0.792797 | 0.469 | 0.469 |
|  | WT-Bf | 35 | 999 | 3.217311 | 0.039 | 0.0585 |
|  | WT-ctrl | 37 | 999 | 3.698057 | 0.024 | 0.048 |
| 3xTg-AD-ctrl | WT-Bf | 30 | 999 | 4.854274 | 0.005 | 0.015 |
|  | WT-ctrl | 32 | 999 | 4.84516 | 0.004 | 0.015 |
| WT-Bf | WT-ctrl | 34 | 999 | 0.854948 | 0.453 | 0.469 |
| Weighted UniFrac pairwise comparisons - amyloidosis and tauopathy |  |  |  |  |  |  |
| Group 1 | Group 2 | Sample size | Permutations | pseudo-F | p-value | q-value |
| 3xTg-AD-Bf | 3xTg-AD-ctrl | 19 | 999 | 1.372471 | 0.236 | 0.372 |
|  | WT-Bf | 17 | 999 | 1.141547 | 0.31 | 0.372 |
|  | WT-ctrl | 16 | 999 | 0.966325 | 0.417 | 0.417 |
| 3xTg-AD-ctrl | WT-Bf | 18 | 999 | 3.94824 | 0.008 | 0.048 |
|  | WT-ctrl | 17 | 999 | 1.258886 | 0.254 | 0.372 |
| WT-Bf | WT-ctrl | 15 | 999 | 1.502895 | 0.195 | 0.372 |

To address the effects of cage and sequencing run on the gut microbiome composition, we performed multivariate PERMANOVA. PERMANOVA was performed at the baseline, amyloidosis, and amyloidosis and tauopathy timepoints. At all time points, cage and sequencing run significantly affected Unweighted UniFrac distance matrices; however, genotype was a more important variable when considering the test statistic (pseudo-F) (Supplemental Tables 10 and 11).

To determine which taxa were differentially abundant across genotype and treatment groups, we performed ANCOM-BC^34^. Results include taxa that were significantly differentially abundant after correction for multiple comparisons and represent features collapsed at Level 7. Taxonomic features are indicated in Fig. 7. At the baseline timepoint, we observed 123 total features across all samples (Fig. 7A-D). At the amyloidosis timepoint, we observed 133 features across all samples. Group comparisons revealed differences in taxa across genotype and treatment type at the amyloidosis timepoint (Fig. 7E-H). At the amyloidosis and tauopathy timepoint, we observed 160 features across all samples. Group comparisons revealed differences in taxa across genotype and treatment type at the amyloidosis and tauopathy timepoint (Fig. 7I-L).

**Figure 7.**
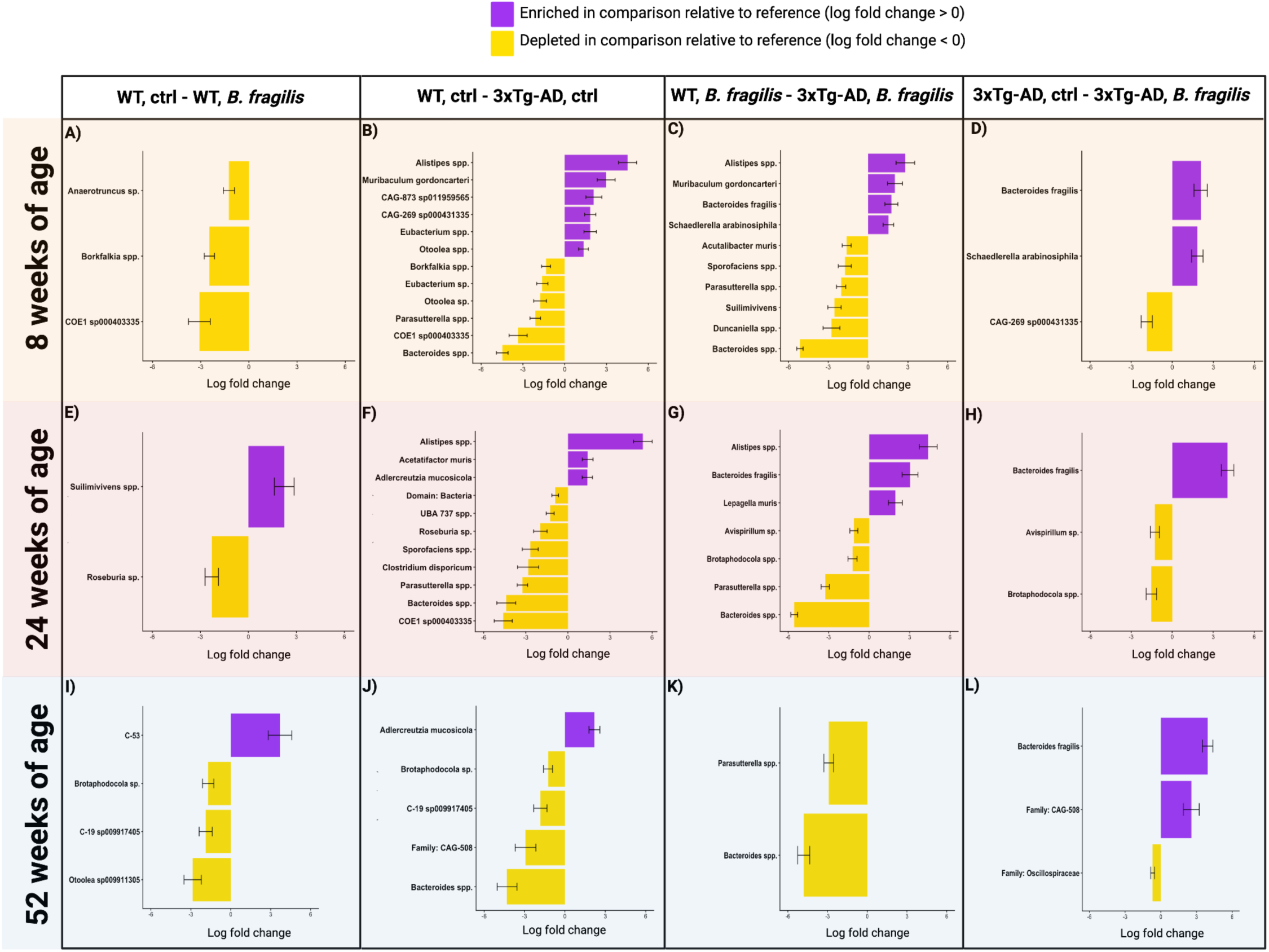
ANCOM-BC log fold change of feature relative abundances between reference and comparison groups (reference-comparison group). Feature log fold changes at eight weeks of age between A) WT mice given control and WT mice given *8. fragi/is,* B) WT mice given control and 3xTg-AD mice given control, C) WT mice given *8. fragi/is* and 3xTg-AD mice given *8. fragilis,* and D) 3xTg-AD mice given control and 3xTg-AD mice given *8. fragilis.* Feature log fold changes at 24 weeks of age between E) WT mice given control and WT mice given *8. fragilis,* F) WT mice given control and 3xTg-AD mice given control, G) WT mice given *8. fragi/is* and 3xTg-AD mice given *8. fragi/is,* and H) 3xTg-AD mice given control and 3xTg-AD mice given *8. fragilis.* Feature log fold changes at 52 weeks of age between I) WT mice given control and WT mice given *8. fragilis,* J) WT mice given control and 3xTg-AD mice given control, K) WT mice given *8. fragilis* and 3xTg-AD mice given *8. fragi/is,* and L) 3xTg-AD mice given control and 3xTg-AD mice given *8. fragilis*.

### *B. fragilis* partially restores metabolite dysregulation in the gut

Differences in fecal metabolome composition at baseline, amyloidosis, and amyloidosis and tauopathy were assessed using PCA of the RCLR-transformed extracted peak areas for both polar and non-polar metabolomes. Analysis on both the polar and non-polar metabolome yielded similar results to the microbiome analysis, with genotype primarily modulating the non-polar metabolome (PERMANOVA: 8 weeks p=0.005, pseudo-F=2.2, n=11; 24 weeks p=0.001, pseudo-F=2.07, n=25; 52-56 weeks p=0.015, pseudo-F=1.79, n=22, Fig. 8A-B). *B. fragilis* treatment shifted the non-polar metabolome, but to a lesser extent (PERMANOVA: genotype-treatment 8 weeks: p=0.001, pseudo-F=1.92, n=11; 24 weeks: 0.001, pseudo-F=1.50, n=25; 52-56 weeks: p=0.001, pseudo-F=1.61, n=22, Fig. 8B; Supplemental Fig. 12C).

**Figure 8.**
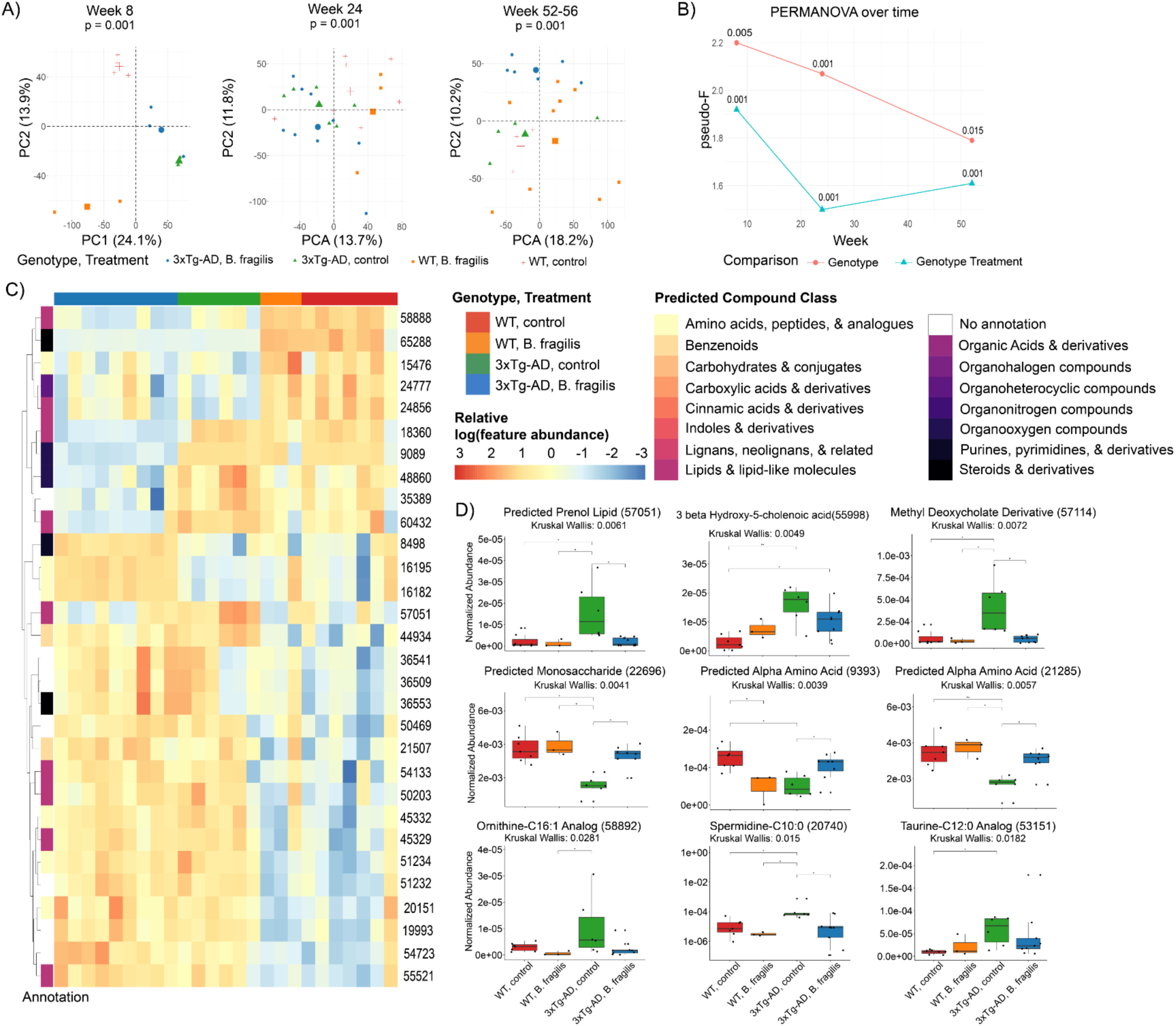
Changes in the fecal non-polar metabolome analyzed using RP UHPLC-MS/MS. A) PCA score plots and PERMANOVA for study weeks 8, 24, and 52-56. Group centroids are enlarged and individual points are colored by genotype-treatment group. B) F-statistic from PERMANOVA calculated at each time point during the study, with calculated p-value indicated above each week. C) Heatmap of significant fecal features at week 24. Each sample is colored based on genotype and treatment type (top bar) and each in silico predicted compound class is differentiated by color on a rainbow scale (side bar).The intensity of color shown in the heatmap is based upon the relative log(feature abundance) for each feature. Each feature ID is indicated on the right side of the heatmap plot. D) Boxplots of a subset of features at week 24. Significance with Kruskal-Wallis p-values for each metabolite are listed, and* indicates p<0.05 in post-hoc Dunn’s analysis with BH correction. Features preceded with “Predicted” (57051, 22696, 9393, and 21285) are annotation level 3 from CANOPUS-predicted compound classes. Other annotations are level 2 annotations from GNPS spectral matching^35^.

Given the observed multivariate separation in our study both by genotype and treatment with *B. fragilis* (Fig. 8A-C, Supplemental Fig. 12A-C), we hypothesized that there would be unique metabolic features differing between genotypes that also differed with *B. fragilis* treatment. To assess this, we performed Kruskal-Wallis and Dunn’s Test analysis on every metabolic feature at 8 weeks of age (n=11 samples), 24 weeks of age (n=25 samples), and 52-56 weeks of age (n=22). (Supplemental Table S12-S18). The top thirty globally differential features for each time point are displayed in clustered heatmaps (Fig. 8C, Supplemental Fig. 13-14), which reveal dysregulation of *in silico* predicted compound classes that include amino acids and derivatives, bile acids and derivatives, and N-acyl lipids (Supplemental Table S19-S30).^30^ At each timepoint, we observed some features which are altered by *B. fragilis* treatment while other features appear unchanged (Supplemental Fig. 15-20, Supplemental Table S22-S24, S28-S30).

Targeted analysis on a subset of neurotransmitters and tryptophan derivatives showed increased tryptophan abundance in 3xTg-AD mice treated with *B. fragilis* as compared to WT treated with *B. fragilis* at week 8, suggesting that *B. fragilis* may alter tryptophan levels in a context-dependent manner (Supplemental Fig. 21) though this dependence is no longer observed at later study timepoints. GABA and glutamine show decreased abundance in *B. fragilis* treated 3xTg-AD mice as compared to vehicle control treated 3xTg-AD mice, though these changes are not statistically significant (Supplemental Fig. 21). Glutamate demonstrated no change across genotype-treatment groups for this 8 week timepoint (Supplemental Fig. 21).

The 24-week (amyloidosis) timepoint had the highest number of significantly differentiated features between WT and vehicle-treated 3xTg-AD mice (Supplemental Table S20, S23). Kruskal-Wallis analysis of polar features at 24 weeks of age revealed that many of the amino acids and *in silico* amino acid derivatives exhibit the same trend as observed in the frontal cortex. Specifically, we observe a decrease in feature abundance in 3xTg-AD mice treated with vehicle control as compared to wild type vehicle control mice that is rescued back to wild type levels after *B. fragilis* treatment in 3xTg-AD mice (Fig. 8C-D). We observe a similar trend in select N-acyl lipids, including a derivative of taurine-C12:0 and spermidine-C10:0 (Fig. 8C-D). The abundance of some bile acids and derivatives followed a similar trend as observed in the N-acyl lipids highlighted above; levels of these metabolites were depleted in *B. fragilis*-treated 3xTg-AD mice (Fig. 8C-D).

Untargeted analysis of fecal samples from the amyloidosis and tauopathy timepoint (52 weeks) revealed fewer differential features between WT and 3xTg-AD vehicle-treated mice and fewer features modulated by both genotype and *B. fragilis* treatment in 3xTg-AD mice Supplemental Tables S21, S24). Included in these features is level 1 annotated cytosine (feature ID: 11254) and a nicotinamide derivative (feature ID: 1171, Supplemental Fig. 20), each of which were modulated towards wild type abundances in *B. fragilis*-treated 3xTg-AD mice. Overall, fecal metabolomics revealed time-dependent genotype- and treatment-associated metabolic alterations, with the most pronounced effects observed early after *B. fragilis* administration, including partial normalization of AD-associated amino acid, bile acid, and N-acyl lipid dysregulation.

## Discussion

In this study, we highlight the critical role of the gut-microbiome-brain axis in AD. We hypothesized that 3xTg-AD mice chronically exposed to *B. fragilis* would demonstrate i) altered gut microbial communities, ii) restoration of neuroinflammation and neurotransmitter activity in the brain, and iii) rescue of metabolites in the brain and gastrointestinal tract to resemble a WT metabolome. We show that 3xTg-AD mice are depleted of native *Bacteroides spp.* which may allow orally-administered *B. fragilis* to colonize the gut microbiome earlier and more consistently than WT mice treated with *B. fragilis*. We also show that *B. fragilis* administration shifts neuroinflammatory gene expression and the brain and fecal metabolomes in 3xTg-AD mice. These results support a mechanistic role of *Bacteroides spp.* observed in some studies of AD^8,9,14,36^.

*B. fragilis* administration altered frontal cortex gene expression of *FOXO3, SLC1A3,* and *GFAP* in a genotype-dependent manner. *FOXO3* is expressed in several types of cells in the brain, including neural cells, astrocytes, and to some extent, microglia^37^. We observed a decrease in *FOXO3* expression in 3xTg-AD mice treated with *B. fragilis*. *FOXO3* has been implicated in several processes related to neuroinflammation and AD pathogenesis. For example, *FOXO3* deficiency in the cortex of aged mice led to aberrant astrocyte activation and reduced amyloid-β uptake capacity^37^. Additional work has demonstrated a role for *FOXO3* in regulating amyloid-β-associated microglial responses^38,39,40,41^. Although microglia contribute to amyloid-β clearance, they may also promote progression of AD pathologies when shifted toward a pro-inflammatory phenotype^42^.

*SLC1A3* encodes the excitatory amino acid transporter 1 (EAAT1), a glutamate transporter expressed primarily on glial cells. EAAT1 transports glutamate from the neuronal synaptic cleft into astrocytes where it reacts with ammonia to form glutamine. *FOXO3* is positively correlated with glutamine synthetase, which converts intracellular glutamate to glutamine^4343–45^. This suggests that the observed correlation between *FOXO3* and *SLC1A3* is potentially important. Together, these findings suggest that B. fragilis may influence astrocyte-associated glutamatergic and neuroinflammatory pathways in mice genetically predisposed to AD pathology.

*GFAP* encodes for a cytoskeletal protein expressed by activated astrocytes and indicates potential metabolic dysfunction in the brain^46^. Increased *GFAP* expression correlates with cognitive decline^47,48^. Increased GFAP can be observed in cognitively normal adults at risk for AD, indicating that glial cell damage or activation precedes cognitive decline^49^. In our study, B. fragilis treatment attenuated the age-associated increase in GFAP expression observed in 3xTg-AD mice, supporting a potentially protective effect on neuroinflammatory processes. Future studies should determine the molecular mechanism underlying *B. fragilis*’ effect on neurotransmitter and neuroinflammatory gene expression.

Our results suggest that *Bacteroides fragilis* mediates fecal microbiota and metabolites, consistent with previous literature^50–53^. *Bacteroides fragilis* altered the gut microbial community composition in a genotype-dependent manner. Consistent with prior studies in mice, we also demonstrated that genotype was a primary driver of gut microbiome composition^8,11,54,55^. Within each genotype, *B. fragilis* modulated microbiome composition. *Bacteroides* taxa have been reported in several studies as enriched or depleted in humans with AD or animals modeling AD pathologies^9,10,14,11,56^. The relative absence of native *Bacteroides* within the 3xTg-AD microbiome may explain the more robust incorporation of *B. fragilis* in 3xTg-AD mice compared to WT controls, as well as the delayed colonization observed in WT mice. Overall, *B. fragilis* treatment reshaped gut microbial community structure in a manner influenced by host genotype and microbial niche availability.

Our results suggests that *B. fragilis* may act as a context-dependent regulator of fecal metabolites involved in neurotransmitter and tryptophan metabolism, though additional follow-up is needed. Prior studies suggest that *Bacteroides* influence tryptophan and neurotransmitter metabolism, with reported alterations in ileal tryptophan metabolites and neurotransmitters^57,17^. Here, serotonin-related metabolites were altered at 24 weeks by *B. fragilis* treatment, including changes in N-acetyl-5-hydroxytryptamine and 3-hydroxybutyroyl tryptamine. *B. fragilis* expresses bile salt hydrolase enzymes that facilitate bile acid deconjugation and formation of secondary bile acids. We observed modulation of bile acid derivatives in *B. fragilis* treatment groups, which were elevated in 3xTg-AD mice relative to WT vehicle-treated and reduced following *B. fragilis* treatment. Altered levels of bile acids have been previously characterized in AD, with some bile acids exhibiting neuroprotective effects and other bile acids exhibiting cytotoxic effects^58^. Beyond bile acids, N-acyl lipids (including ornithine-, spermidine-, and taurine-conjugates) were altered by *B. fragilis* treatment, consistent with previous studies in AD mouse models^59^ and patient-derived datasets^60^. While limited information is available regarding this relationship, modulation of N-acyl lipids in the gut has been linked to differences in microbial colonization^61^.

Treatment with *B. fragilis* shifted the frontal cortex metabolome in 3xTg-AD mice towards that of the WT controls at the amyloidosis and tauopathy timepoint. The frontal cortex metabolome may be influenced via the interaction of *Bacteroides* with the host, as the 3xTg-AD mice given vehicle control are the only group in this study which don’t host a native or supplemented *Bacteroides.* Amino acids and amino acid-derived metabolites represented a major class distinguishing WT and 3xTg-AD mice. Treatment with *B. fragilis* was associated with a partial normalization of several amino acid derivatives in 3xTg-AD mice to WT levels. This may reflect direct microbial effects on amino acid metabolism or indirect effects mediated by changes in host physiology consistent with improved health status. Dysregulation of amino acids have been widely reported in AD patients^62^, transgenic mice modeling AD^63,64^, and mice receiving FMT from AD patients^65^, with dietary interventions targeting certain amino acid pathways showing therapeutic potential^66–69^. Notably, commensal *B. fragilis* secretes amino acids, including glutamate, tryptophan, and ornithine, which may contribute to these changes observed post-treatment^50^. As with amino acid metabolism, both pyrimidine metabolism and nicotinamide-dependent pathways are dysregulated in AD^70,71^ and have been explored as therapeutic targets ^72–74,71,75^. At 52-56 weeks, cytosine and a nicotinamide-derivative were modulated by *B. fragilis*. Further investigation is needed to determine the mechanisms underlying these associations.

Results from this study suggest that chronic exposure to *B. fragilis* affects broad metabolic, glutamatergic, and potentially GABAergic signaling pathways in mice modeling AD pathologies. Further research is needed to establish the mechanisms underlying this communication and how genetic predisposition to develop the pathologies of AD interacts with gut microbiome-derived metabolic byproducts and gliosis.

### Limitations and future directions

We acknowledge that our study employs a mouse model of AD pathologies and that any use of a model organism may not translate into humans. We selected female mice in this study because females experience a higher incidence of AD^76^ and female 3xTg-AD mice express the pathologies of AD more consistently than male mice^77,78^. Mice in this study were aged until they expressed the hallmark pathologies of AD, which does not represent a physiologically “elderly” time point. Aging mice to 18 or 24 months may demonstrate a greater effect of disease pathogenesis. We assessed cognitive outcomes via the Morris Water Maze and Novel Object Recognition task at 52-56 weeks but did not observe any significant differences across genotypes or experimental groups (data not shown). Since we observed the most robust differences in the microbiome earlier in life, we will include the cognitive battery at earlier timepoints, such as when amyloidosis, but not tauopathy, is modeled. We carefully considered housing strategies, and we chose to house genotypes separately and reduce mouse density per cage to include multiple cages per experimental group as suggested by Kim et al^79^. We selected not to co-house 3xTg-AD mice with WT mice because several studies have described a partial transfer of AD microbiota, increased neuroinflammation, and behavioral deficits in WT mice co-housed with models of AD pathologies^54,80,81^. As expected, we observed cage effects but these were not the primary drivers of the observed compositional changes in the gut microbiome, rather we observed stronger genotype-mediated effects.

## Acknowledgements

We would like to thank the Biological Sciences Vivarium staff at Northern Arizona University for animal husbandry, especially Kathleen Freel and Kimberly Cohen.

This project was funded by the Arizona Alzheimer’s Consortium (AAC, PI: Cope) and NIH R21 (1R21AG074203, PI: Cope). K.A.C was supported through the Arizona Technology Research Initiative Funds (TRIF). A.T.A and O.S. were also supported by funds from the Boettcher Foundation’s Webb-Waring Biomedical Research Program.

